# Sex and experience dependent regulation of synaptic protein turnover

**DOI:** 10.1101/2025.11.24.690161

**Authors:** Seok Heo, Shiyu Zhang, Dong-Gi Mun, Akhilesh Pandey, Alexei M Bygrave, Richard L Huganir

## Abstract

Synaptic transmission can be carefully tuned through plasticity mechanisms that regulate synaptic strength, structure, and number. In vivo measurements demonstrate remarkable spine dynamics, with subsets of synapses persisting for months. This correlates with the extreme longevity of certain memories, which can persist for an organism’s lifetime. The molecular basis supporting the long-term stability of specific synapses and the long-term durability of memories remains unknown. At the protein level, most proteins persist for a relatively short amount of time before they are degraded and replaced with new molecules. However, recent work has identified a population of proteins, including those present at the synapse, that are exceptionally long-lived. It has been speculated that long-lived proteins (LLPs) could contribute to long-term synapse stability, function, and memory. Here, we used stable isotope labeling in mammals (SILAM) to first identify LLPs in the post synaptic density (PSD) of the hippocampus and subsequently determine if protein turnover rates varied by sex or following learning. We identified novel synaptic LLPs and found that both sex and experience can regulate synaptic protein turnover rates. We identified sex-dependent changes in protein turnover rates in autism spectrum disorder (ASD) risk genes, including increased stability of Gabrg2, a GABA-A receptor subunit, in male mice. Furthermore, we observed stabilization of a subset of PSD proteins, such as Shank3, following contextual fear conditioning. We propose that sex and experience dependent changes in protein turnover rates could help explain sex-differences in psychiatric risk and aid our understanding of the molecular mechanisms that support learning and memory.

## Introduction

Individual proteins have finely tuned rates of synthesis and degradation, with lifetimes spanning several orders of magnitude. Post mitotic cells, including neurons, have been shown to contain a subset of particularly stable proteins, often referred to as long-lived proteins (LLPs). In brain tissue, examples of LLPs include nuclear pore complex proteins (Savas et al., 2012), components of the extracellular matrix, as well as documented synaptic proteins (Fornasiero et al., 2018; Heo et al., 2018; Li et al., 2025). Presently, the contribution of LLPs to synapse function remains enigmatic.

Neuronal cells have some unique challenges to overcome as they typically exhibit long-range axonal projections and extensive dendritic arborizations, for which they need to maintain integrity and proper function for the lifetime of the organism. With the development of *in vivo* imaging methodologies, it has been shown that individual synapses have huge variability in their longevity, with some very transient and others persisting for months (Trachtenberg et al., 2002). The stability of some synapses is quite remarkable and readily exceeds the estimated turnover rate of most synaptic proteins. With synaptic plasticity thought to be a correlate of learning and memory a paradox emerges: how can changes in synapse function (i.e. LTP and spine dynamics) be maintained longer than the lifetimes of the proteins that form the constituent parts of the synapse? The synaptic tagging hypothesis posits that activated synapses contain a biomolecular tag that promotes subsequent—and long term—sequestration of proteins which support LTP (Frey and Morris, 1998). LLPs have been identified at the synapse (Heo et al., 2018), raising the question as to whether a subset of exceptionally stable proteins could represent a “synaptic tag” to support the long-term stability of specific synapses, or even the maintenance of long-term memories.

Protein turnover rates are frequently assessed using stable isotope labeling in mammals (SILAM) coupled with mass spectrometry, wherein protein degradation and subsequent synthesis lead to changes in the relative abundance of heavy isotopes within individual proteins. Previous research shows that protein turnover rates are dependent on cellular environment as the presence of Glia has been shown to increase the turnover rates of neuronal proteins *in vitro* (Dörrbaum et al., 2018). Additionally, the experience of environmental enrichment has been shown to alter protein turnover rates of synaptic proteins *in vivo* (Fornasiero et al., 2018; Heo et al., 2018). Currently, whether protein turnover rates are altered by sex remains unknown. This is a pressing issue because there are well documented sex differences in the prevalence of psychiatric disorders. For example, autism spectrum disorder (ASD) is more readily diagnosed in males (Loomes et al., 2017). While acknowledging that this could also be attributed to missed diagnosis in women, it also suggests possible biological sex differences underlying ASD. SFARI has collated ASD risk genes (https://gene.sfari.org/), many of which are established synaptic proteins.

In the present study, we sought to document how turnover rates of the hippocampal synaptic proteome changes with sex, or after a fear-learning behavioral task. Our analysis focused on the postsynaptic density (PSD) of the hippocampus. As expected, we identified a subset of proteins with long half-lives. Curiously, we found sex differences in synaptic protein turnover includes many autism spectrum disorder risk genes. Additionally, we show that a subset of synaptic proteins tune their turnover rates following contextual fear conditioning.

## Results

### Identification of stable synaptic proteins

We conducted a stable isotope labeling in mammals (SILAM) experiment to characterize the relative turnover rates of proteins enriched at the postsynaptic density (PSD). To promote maximal isotope labeling, chow containing Lys-^13^C_6_ was given to a breeding pair so labeling could begin *in utero*. When the resulting offspring reached 10 weeks, they were switched to a control diet with regular Lys-^12^C_6_ to initiate a “chase” period (**Figure 1A**). Tissue was collected at different time points and biochemical fractionation performed on hippocampal tissue to extract the postsynaptic density (PSD) fraction (**Figure 1B**). At time point 0 the average relative isotope abundance (RIA) for each protein was 0.87 for the PSD fraction, indicating very high levels of labeling, with 2961 proteins having a starting RIA >0.8 (**Figure 1C**).

**Figure 1.**
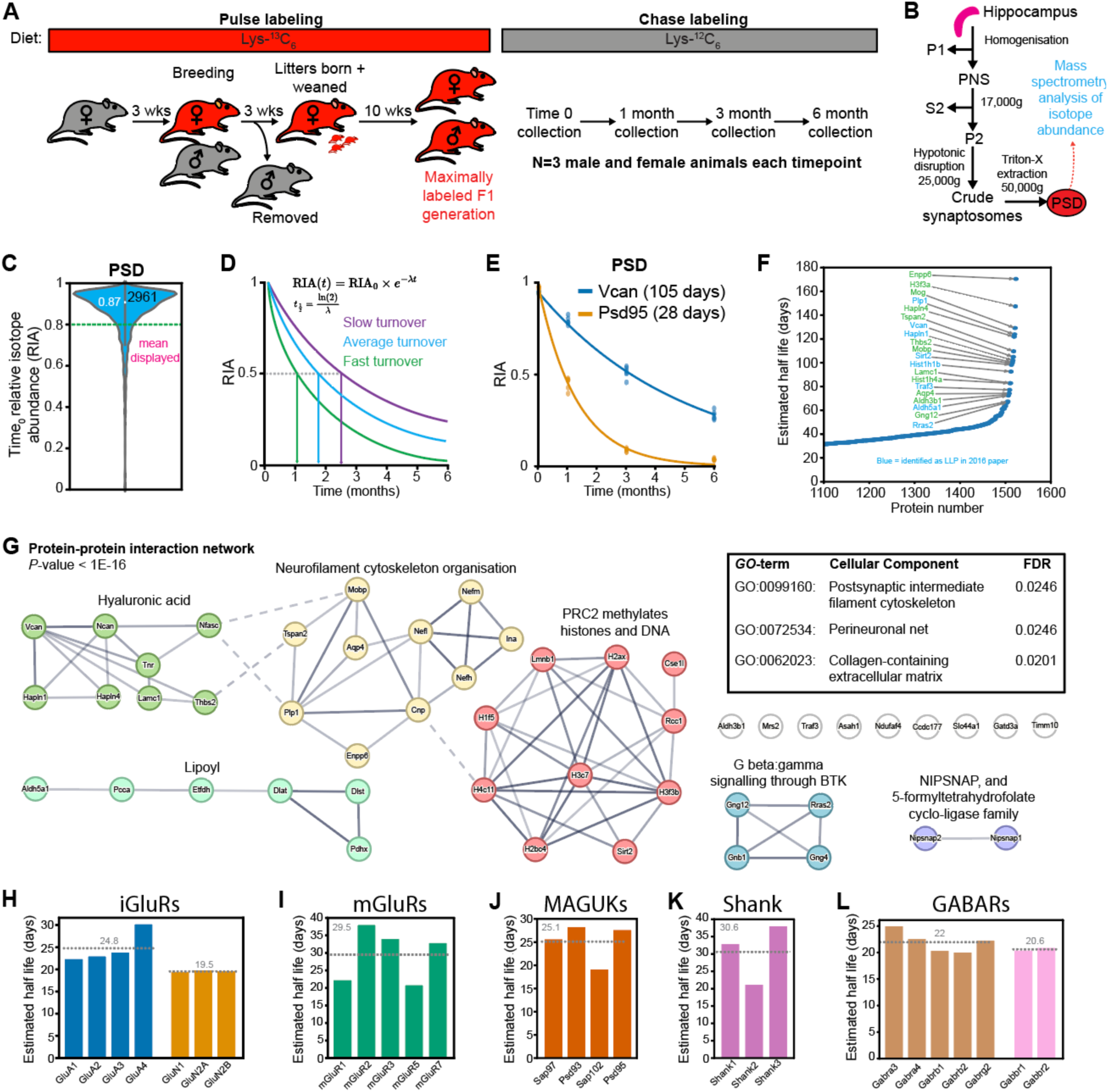
Identification of stable synaptic proteins. (**A**) Schematic of pulse chase SILAM labeling scheme. (**B**) Schematic of tissue processing before mass spectrometry. (**C**) Violin plot displaying the relative isotope abundance (RIA) values for each protein at the beginning of the chase period (Time0). The green dotted line shows the threshold for inclusion (>0.8) and the white point displays the mean (0.87). (**D**) Schematic showing fit of exponential decay of the RIA values over time, with examples of curves for slow, average and fast turnover. (**E**) Data showing example of proteins with average (Psd95, 28 days) or stable (Vcan, 105 days) estimated half-lives. Each data point represents an individual animal. (**F**) Scatter plot of individual proteins arranged by relative stability. The top 30 proteins are labelled. Blue labels indicate proteins we previously described as stable synaptic proteins (Heo et al., 2018). (**G**) STRING analysis of the top 50 stable proteins. Proteins are colored by their associated cluster labels. A significant protein-protein interaction network indicates that proteins are more connected than would be expected by chance. Inset box shows *GO*-terms for cellular components that were significantly enriched in these 50 proteins compared to a background of all proteins in our PSD dataset. (**H-L**) manual curation of the estimated half-lives of proteins in key receptor or scaffolding protein families. The gray dotted line indicates the average turnover rate for each group.

Previously, Fornasiero *et al*. measured the stability of proteins in the brain by implementing a double exponential function that considers the recycled pool of amino acids containing the heavy isotope, providing a precise calculation of protein lifetime (Fornasiero et al., 2018). We applied this method to our data, using their reported constant(s). Unexpectedly, we observed that many proteins in our dataset had estimated half-lives with values close to zero (**Supplementary Figure 1A**). Of note, the constant(s) used by Fornasiero *et al*., were derived from male mice only. To see if sex impacted our implementation of their formula, we split our data into male and female animals. There was a larger proportion of close-to-zero protein half-lives within the isolated female dataset (**Supplementary Figure 1B**). Because the constant used in this calculation method is acutely sensitive to differences in metabolism rates likely different in male and female mice, a separate calibration experiment would be required to calculate appropriate constants for female animals. Indeed, it is likely that any experimental group where metabolism could be different will require its own calibration experiments.

Therefore, to circumvent this confound, we proceeded by evaluating turnover rates using a simple exponential decay function. At each time point during the chase period the RIA was calculated for each protein. By fitting an exponential decay function to the RIA values, we estimated the half-life of each protein, where slower RIA decay indicates proteins undergo slower degradation (**Figure 1D**). This revealed proteins with vastly different turnover times, exemplified by Vcan and Psd95 (**Figure 1E**). Reassuringly, we observed a strong positive and highly significant correlation between protein half-lives in our PSD dataset (estimated by simple exponential decay) and half-lives reported by Fornasiero *et al*., in the synaptosome fraction (**Supplementary Figure 1C**). We acknowledge that we are likely overestimating protein half-lives and therefore focus on the *relative* turnover rates between proteins.

We identified a subset of PSD proteins with exceptionally slow relative turnover rates, including proteins found in our previous study such as Vcan, Traf3 and Rras2 (Heo et al., 2018), as well as novel proteins (**Figure 1F**; **Supplementary Table 1**). We conducted a STRING analysis of the top 50 most stable proteins. These proteins were significantly connected and were enriched for *GO*-terms such as postsynaptic intermediate filament cytoskeleton, perineuronal net, and collagen-containing extracellular matrix (**Figure 1G**). We conducted a manual evaluation of relative half-lives of different ionotropic glutamate receptors (**Figure 1H**), metabotropic glutamate receptors (**Figure 1I**), membrane-associated guanylate kinase family members (**Figure 1J**), SH3 and multiple ankyrin repeat domain containing proteins (**Figure 1K**), and GABA receptors (**Figure 1L**). It was interesting to observe differences in the half-lives of proteins in the same family, such as the relative stability of GluA4 (**Figure 1H**), an AMPA receptor subunit particularly enriched in GABAergic inhibitory interneurons (Geiger et al., 1995), as well as the reduced stability of Sap102 (**Figure 1J**), a protein known to be more mobile at synapses compared to other MAGUKs (Zheng et al., 2010).

### Sex differences in protein turnover

To our knowledge, previous studies looking at protein turnover rates have not considered sex differences. We were interested to probe for basal changes in protein turnover based on sex, as there are well documented sex differences in the prevenance of psychiatric disease (Yang et al., 2024) and neurological disorders (Bonkhoff et al., 2025). Therefore, we split our dataset into male and female animals and recalculated relative protein turnover rates. When analyzed separately there was no difference in the initial RIA values in male and female mice, indicating that incorporation of the heavy isotope was not sex dependent (**Supplementary Figure 2A, B**). We observed a strong correlation between protein stability in male and female mice (**Figure 2A**). Interestingly, there was an overall trend towards proteins being more stable in male mice. Differences in metabolism likely underlie this general trend. In fact, female mice consumed significantly less food in both the labelling and chase phases (**Supplementary Figure 2C, D**). A subset of proteins appeared to be differentially stable in male and female mice. We considered proteins differentially regulated by sex as those that had a male/female half-life ratio of greater or less than 2 standard deviations of the mean (**Figure 2A; Supplementary Table 2)**. We noticed that several proteins exhibiting sex differences in their stability were also autism spectrum disorder (ASD) risk genes, as designated by the SAFARI database (https://gene.sfari.org/). STRING analysis of these proteins with sex differences revealed significantly connected proteins, with clusters relating to inhibitory synapses and signal transduction by L1 (**Figure 2B**).

**Figure 2.**
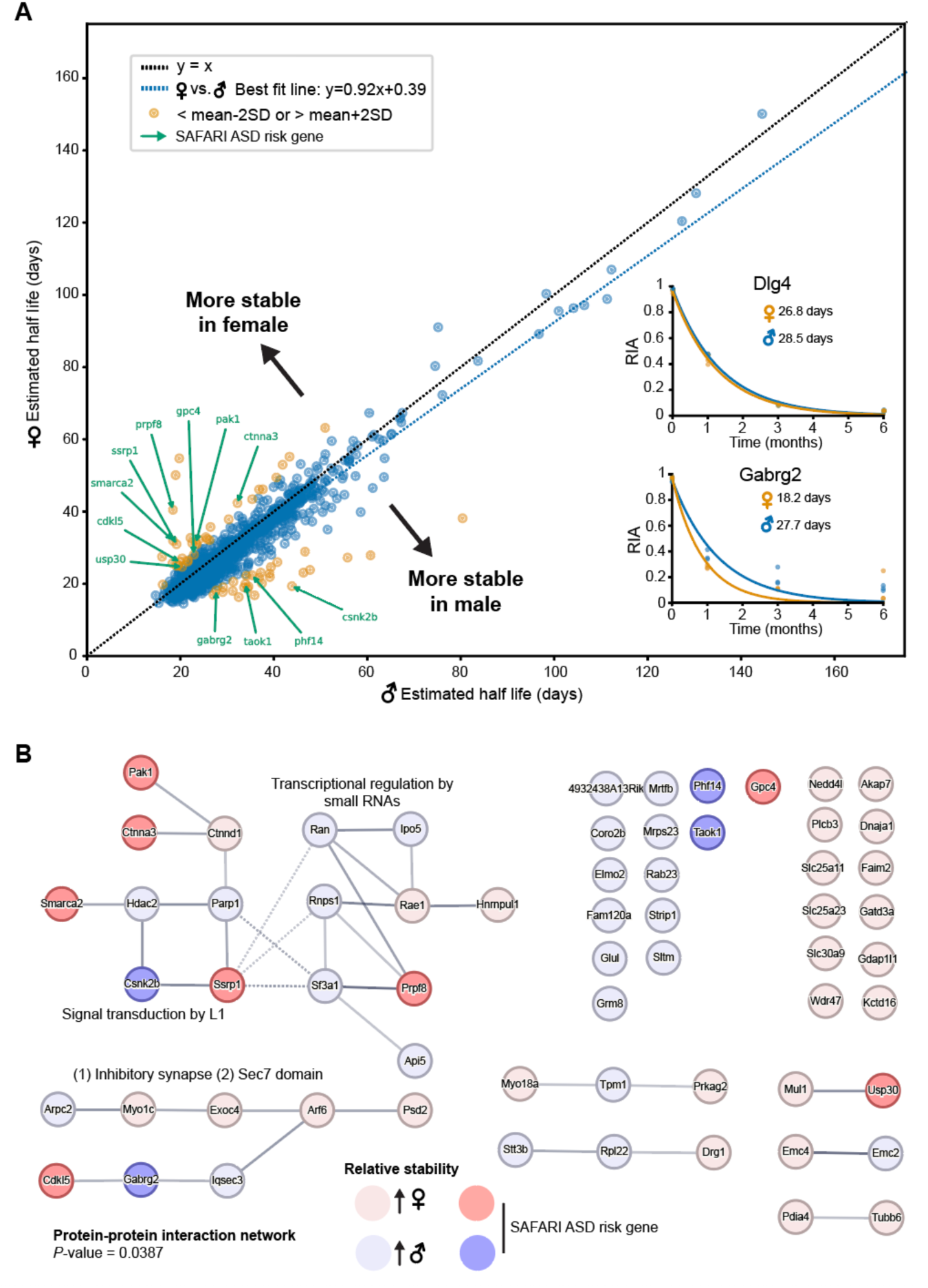
Sex differences in protein turnover. (**A**) Scatter plot showing the relative turnover rates of individual proteins in male (x-axis) and female (y-axis) animals. Yellow symbols indicate proteins that were +/− 2 standard deviations (SD) from the mean. Green arrows and labels identify putative autism spectrum disorder (ASD) risk genes from the SAFARI database. Inset graphs indicate raw data from two example proteins that do not change with sex (Dlg4) or that show greater stability in male mice (Gabrg2). Note, the black dotted line indicates unity between the axes. The blue dotted line shows the best fit of the data, highlighting that proteins were, in general, slightly more stable in male animals. (**B**) STRING analysis of the proteins with +/− 2 SD differences in turnover rates. Blue shading indicates greater stability in male animals and red shading indicates greater stability in female mice. Darker shading in blue or red indicates putative ASD risk genes.

### Aversive experience tunes stability of synaptic proteins

We next sought to assess changes in hippocampal PSD protein stability following contextual fear conditioning (cFC). cFC is a well-established hippocampus-dependent task that triggers long term memory following a single training session (**Figure 3A**). We confirmed that contextual fear conditioning induces long-term memory in a pilot study with unlabeled animals, as demonstrated by freezing behavior when returned to the conditioning context. In this cohort, aversive memories persisting up to two months compared to control animals that received no foot shock in the arena (**Figure 3B**).

**Figure 3.**
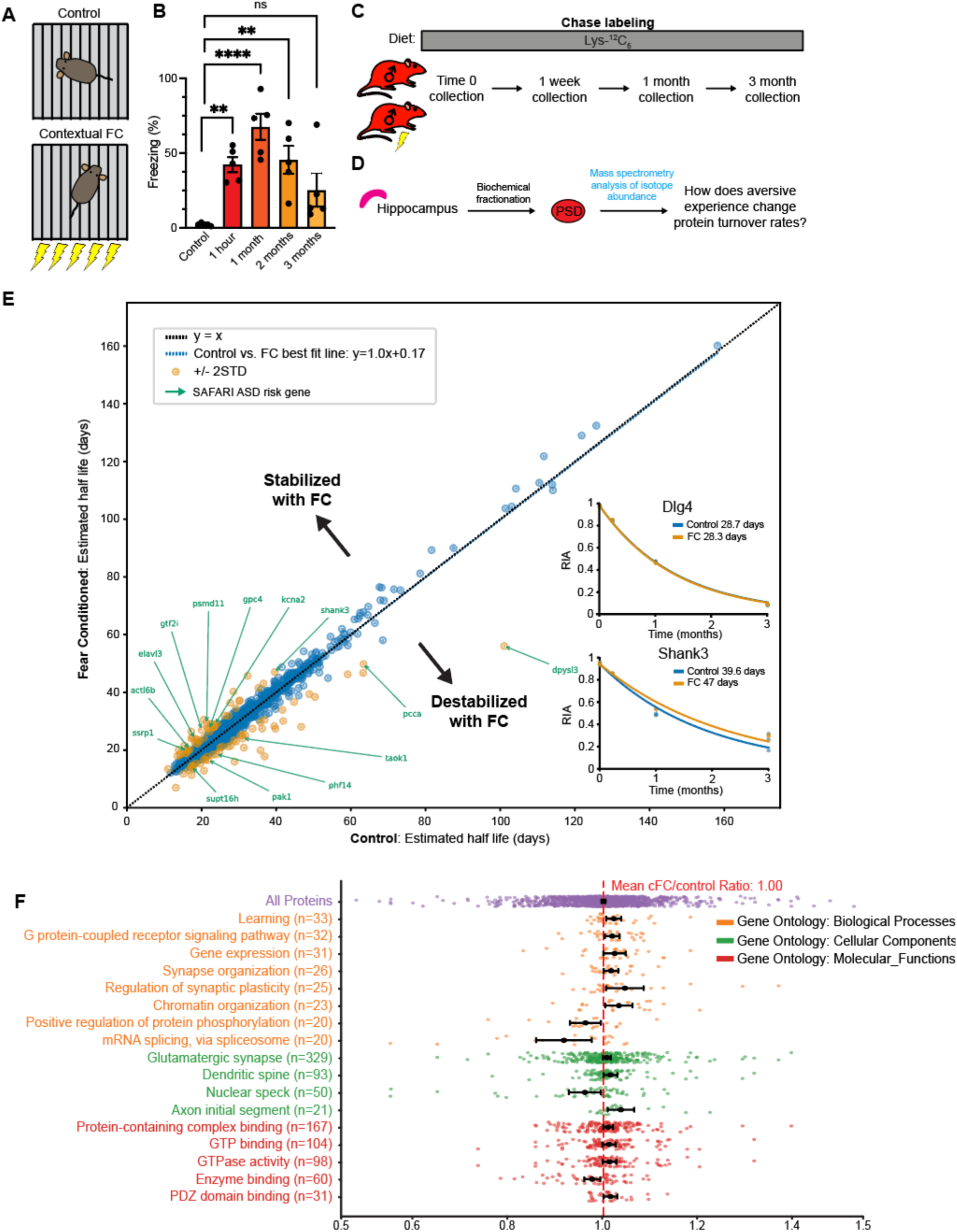
Aversive experience tunes stability of synaptic proteins. (**A**) Schematic of the contextual fear conditioning (cFC) paradigm. (**B**) Pilot study in unlabeled mice demonstrating robust freezing upon re-exposure to the context, used as a proxy for the fear memory. All groups were compared to the control group, which did not receive foot shocks while in the apparatus. (**C**) Labeling strategy and time course for tissue collection. Note, in this experiment only male mice were included. (**D**) Hippocampal tissues was subject to biochemical fractionation and subsequent mass spectrometry to evaluate how experience changes long-term protein stability. (**E**) Scatter plot showing the relative turnover rates of individual proteins in control (x-axis) and cFC (y-axis) animals. Yellow symbols indicate proteins that were +/− 2 standard deviations (SD) from the mean. Green arrows and labels identify putative autism spectrum disorder (ASD) risk genes from the SAFARI database. Inset graphs indicate raw data from two example proteins that do not change with experience (Dlg4) or that show greater stability following cFC (Shank3). Note, the black dotted line indicates unity between the axes. The blue dotted line shows the best fit of the data, highlighting that protein turnover rates were, in general, unchanged with cFC. (**F**) Plot showing the relative change in turnover rates (cFC/control) for all proteins (purple), or proteins linked to Gene Ontology biological processes (orange), cellular components (green) or molecular functions (red) that were significantly different to a ratio of 1 (one-sample T-test) and have 20 or more protein under the GO terms (for statistical details see **Supplementary Table 4**) Error bars display the standard error of the mean. The red dotted line indicates no change in turnover rates with cFC.

Subsequently, we repeated this protocol with labelled male assigned to control or cFC groups. On the conditioning day, animals assigned to the cFC group began freezing after foot shocks were delivered (**Supplementary Figure 3A**). Tissue was collected from control or cFC mice at various timepoints: immediately after training (to serve as the baseline), or following re-exposure to the area at 1 week, 1 month, or 3 months. The PSD fraction was then isolated and subject to mass spectrometry to determine the RIA of each protein (**Figure 3C,D**). As describe above, we fit an exponential decay to the RIA values measured for each protein over time to estimate the relative half-lives (as in **Figure 1D**). The turnover rate of most proteins did not change following FC. This suggests that there are no global changes in protein turnover, or changes in metabolism. Supporting this, the food intake was not changed in the “chase” period in control and FC animals (**Supplementary Figure 3C**). However, there were some exceptions that appeared to be differentially regulated following FC (**Figure 3E**). We considered proteins differentially regulated if the FC/control half-life ratio was greater or less than 2 standard deviations of the mean (**Figure 3E; Supplementary Table 3**). STRING analysis of these differentially regulated proteins revealed a significant protein-protein interaction network, with clusters of proteins related to regulation of dendritic spine morphogenesis and CRMPs in Sema3A signaling (**Supplementary Figure 3D**).

Subsequently, to evaluate potentially subtle changes in turnover rates of related proteins we examined the FC/control turnover ratios of proteins in our dataset linked to specific Gene Ontology biological processes, cellular components, and molecular functions (**Figure 3F**). We only included terms when our dataset included >20 proteins and conducted one-sample T-tests against a FC/control ratio of 1 (i.e. no change). Subtle changes were observed, with groups of proteins related to “learning”, “synaptic organization”, “dendritic spine”, and “PDZ-domain binding” showing elevated stability following FC, and terms such as “positive regulation of protein phosphorylation” showing decreased stability (**Figure 3F**). Curiously, there was no correlation between overall protein stability and the FC/control ratio (**Supplementary Figure 3E**).

## Discussion

Unlike other biomolecules that exhibit longevity, such as DNA, proteins do not have the same repair mechanisms. LLPs are therefore predicted to be particularly vulnerable to accumulating oxidative damage over time, posing an inherent risk to cells that maintain LLPs. Consequently, we hypothesized that LLPs must serve an important function to balance out the risk that accumulation of damaged proteins could impart. The goal of this study was to first identify novel LLPs, and then to assess sex and experience dependence changes in protein turnover rate.

Presently, the role of exceptionally stable LLPs remains largely unknown but could explain how certain synapses exhibit remarkable stability *in vivo*. We used an unbiased approach to evaluate the relative turnover rates of proteins enriched in the PSD, a subcellular compartment essential for synaptic transmission and synaptic plasticity. As expected, based on related studies (Fornasiero et al., 2018; Heo et al., 2018), we identified a subset of particularly stable PSD synaptic proteins. several of the LLPs we identified were extracellular matrix proteins such as Versican, Neurocan, and hyaluronan and proteoglycan link proteins 1 & 4. While not components of the PSD, we expect these proteins are in close association with the PSD and remain in complex throughout the PSD fractionation protocol. Within synaptic protein families we observed differences in the turnover rates of individual proteins. For example, of the AMPA-type glutamate receptors, the GluA4 subunit which is particularly enriched in GABAergic inhibitory interneurons (Geiger et al., 1997; Pelkey et al., 2015) exhibited the slowest relative turnover rate. Parvalbumin positive inhibitory interneurons have been shown to have more stable glutamatergic synapses than their counterparts in excitatory neurons (Melander et al., 2021). It will be fascinating to uncover the molecular basis for the cell type-specific regulation of synapse stability, and to determine if LLPs contribute to such cellular specializations. Within the MAGUK family, Sap102 had the fastest relative turnover rate and is also known to be substantially more mobile at glutamatergic synapses (Zheng et al., 2010). However, we acknowledge that the overall stability of a protein need not match its mobility within a neuron. A PSD protein that could be of particular interest is TNF alpha receptor associated factor 3 (Traf3). A role for Traf3 at the synapse has not been documented, but it has been identified in Psd95 protein complexes previously (Fernández et al., 2009). Furthermore, we recently identified Traf3 as being enriched in Psd95 complexes originating from GABAergic inhibitory interneurons (Bygrave et al., 2023). Interestingly, recent GWAS have linked Traf3 to major depressive disorder (Levey et al., 2021), suicide (Kimbrel et al., 2023), and post-traumatic stress disorder (Nievergelt et al., 2024). How the stability of Traf3 could relate to its function at the synapse remains to be investigated.

Sex differences in the prevenance of psychiatric disorders are well documented. For example, ASD is much more readily diagnosed in males (Loomes et al., 2017), and major depressive disorder is more prevalent in females (Cyranowski et al., 2000). Previous studies evaluating protein turnover rates have not considered sex as a variable. In the present study, we identified a subset of proteins that were differentially stable in male and female mice. Curiously, a number of these proteins were putative ASD risk genes, such as Gabrg2 and Cdkl5. We speculate that sex differences in protein stability—if also present in humans—could contribute to sex-differences in the prevalence of psychiatric disorders. More work is required in this area, but our work highlights the necessity to consider sex-differences when evaluating protein turnover.

A major goal of our study was to identify proteins whose turnover rates change following learning. We picked contextual fear conditioning (cFC) as a learning paradigm at this task produces robust one-trail learning. Overall, the average turnover rate of PSD proteins did not change with cFC, but there were subsets of proteins that were stabilized after cFC, such as Shank3. Shank3 is well documented to play a role in synaptic plasticity and learning and memory (Wang et al., 2011). Therefore, stabilizing Shank3 following cFC could serve as a potential mechanism to promote long-term synaptic potentiation, supporting learning. However, we would like to highlight a caveat to this interpretation as we are currently unable to disambiguate between “learning-dependent” vs. “experience dependent” changes in protein turnover. For example, it is possible that some of the changes we observed are a result of the aversive experience and stress induced by the foot shocks rather than learning *per se*. Future studies could include a shock only group to help to account for this possibility.

With respect to the molecular basis of long-term memory, particular attention has been given to extracellular matrix proteins that form perineuronal nets around neurons and synapses as it has been postulated that changes in perineuronal nets could serve as a stable scaffold supporting long term memory (Tsien, 2013). While composed of very stable proteins, the ECM retains the ability for highly dynamic changes, creating time windows permissive for synaptic plasticity (Dankovich and Rizzoli, 2022). In the present study we did not observe experience dependent changes in ECM protein turnover rates that might have been expected based on the role of ECM remodeling in learning and memory. However, it is possible that evaluating the entire hippocampus masks subtle changes that could be restricted to specific inputs or synapse populations. Recently a new approach has been utilized to track the stability of individual proteins over time using HaloTag-labeled endogenous proteins in conjunction with photostable dyes. This enables time locked labeling of proteins with distinct fluorescent dyes to map the turnover rates while also preserving spatial information. For example, this has been done with GluA2 and Psd-95 (Bulovaite et al., 2022; Mohar et al., 2025). This has shown that in older mice, Psd-95 turnover is slower (Bulovaite et al., 2022), and that the ionotropic glutamate receptor GluA2 shows region specific increases in turnover following acquisition of a new rule in a spatial learning task (Mohar et al., 2025). In the future, using halo-tagged KI mice for specific ECM proteins—and other candidate proteins—in pulse/chase experiments could provide a sensitive way to map how protein turnover could be regulated in a manner to support synaptic plasticity.

Limitations of the study: Our approach was tailored towards detecting proteins with slow rather than fast turnover rates. A modified experiment design with closer spaced timepoints would be useful in evaluating proteins with faster turnover rates, which could also play important roles in regulating synapse function. Finally, our approach lacks the spatial resolution for where changes in protein turnover rates are occurring. As a result, we recognize that if turnover rates change in opposite directions in different subregions of the hippocampus, then these changes could be averaged out and missed in our analysis. This information could be determined with generation of HaloTag-KI proteins and sequential labelling with fluorescent dyes.

## Methods

### Animals

C57Bl/6J animals age were purchased from (Jackson Laboratory) and either used to establish a behavioral paradigm that induces long-term learning and to set up breeding pairs for the stable isotope labeling in mammals experiment. Animal care, use, and experimental protocols were approved by the Institutional Animal and Use Committee (IACUC) of Johns Hopkins University.

### Stable isotope labeling in mammals (SILAM)

Female C57Bl/6J mice were switched to a “heavy diet” (Cambridge Isotope Laboratories, Lys-^13^C6) for 3 weeks before being paired up with male C57Bl/6J animals. Male animals were removed when females were visibly pregnant, or at 3 weeks. When litters were born, and after weaning, mice remained on the “heavy diet” until 10 weeks old for the pulse labeling phase, at which point tissue was collected from a subset of the mice, and food switched to a “light diet” (Cambridge Isotope Laboratories, Lys-^12^C6) for the chase labeling phase. When 10 weeks the animals were allocated into cohorts for two experiments, which were run at the same time:

#### Experiment 1: Sex specific changes in protein stability

Tissue from male and female animals (n=3-4 per sex/time point) was collected at 0, 1, 3, and 6 months following the switch to “light diet”. Data from these groups was used to identify long-lived proteins and to evaluate sex differences in protein turnover. Note: all animals in experiment 1 experienced a brief exposure to the fear conditioning chamber at time point 0, with no shock delivery.

#### Experiment 2: Experience dependent changes in protein stability

An additional group of male-only animals experienced contextual fear conditioning (FC) at time point 0 (details below). Tissue from these mice was then collected at 1 week, 1 month, 3 months, and 6 months. 3 male mice were allocated to a control group that were exposed to the fear conditioning chamber but did not receive any shocks. These animals were also collected 1 week after the chase labelling began. We included the 1-week time point for experiment 2 to help capture early changes in protein stability following the aversive experience of fear conditioning that might be missed using only the 0, 1, 3, and 6 month time points. Data from male animals in experiment 1 was used as a control group for experiment 2.

### Contextual fear conditioning

Ahead of testing mice were transitioned to a reverse light cycle room and all behavioral testing was conducted during the dark (active phase). Mice were habituated with gentle handling for the days preceding testing. Animals were placed into fear conditioning chambers for 300 seconds and given 120s to acclimatize to the arena. Mice in the fear conditioning group (FC) received 5 foot shocks (0.75mA, 2s) delivered at 120s, 150s, 180s, 210s and 240s and were returned to their home cage. Control mice (Con) were placed in the same arena for 300s but no shocks were delivered. This provided a control for context exposure. At different timepoints, animals were re-exposed to the same arena and their freezing was measured over the course of a 300s recall trial. Freezing was considered as a measure for learning.

#### Establishing the task

Male C57Bl/6J mice aged 10 weeks were allocated into different groups: Con, 1hr FC, 1 month FC, 2 months FC and 3 months FC. 5 mice were present in each group. Control group data was from the recall session at 1 hr.

#### During the SILAM experiment

SILAM labeled male mice were assigned into Con or FC groups. Tissue was collected at time point 0 (4 mice, 2 Con, 2 FC) or immediately following the recall session after 1 week (6 mice, 3 Con, 3 FC), 1 month (6 mice, 3 Con, 3 FC), 3 months (6 mice, 3 Con, 3 FC), and 6 months (6 mice, 3 Con, 3 FC).

### Biochemical fractionation

At the respective timepoints the hippocampus was rapidly dissected, and flash frozen in liquid nitrogen. Samples were stored at −80C. The hippocampus from each mouse was homogenized individually using 20 strokes from syringes equipped with 26G x 3/8 (0.45mm × 10mm) needles in homogenization buffer (320 mM sucrose, 5 mM sodium pyrophosphate, 1 mM EDTA, 10mM HEPES pH 7.4, 200 nM okadaic acid, 1 mM sodium orthovanadate, protease inhibitor cocktail (Roche), phosphatase inhibitor cocktail (Sigma-Aldrich)). The homogenate was then centrifuged at 1,000 x g for 10 minutes at 4°C to yield P1 (nuclear fraction) and post-nuclear supernatant (PNS) fractions. PNS fraction was further centrifuged at 17,000 x g for 20 minutes at 4°C to yield P2 (membrane/crude synaptosome) and S2 (cytosol) fractions. P2 was resuspended in hypotonic resuspension buffer (Milli-Q® water with 5 mM sodium pyrophosphate, 1 mM EDTA, 10mM HEPES pH 7.4, 200 nM okadaic acid, 1 mM sodium orthovanadate, protease inhibitor cocktail (Roche), phosphatase inhibitor cocktail Roche)), then centrifuged at 25,000 x g for 20 minutes at 4°C to yield lysed synaptosome (LS) fractions. Collected LS fractions were resuspended in resuspension buffer (50 mM HEPES pH 7.4, 5 mM sodium pyrophosphate, 1 mM EDTA, 200 nM okadaic acid, 1 mM sodium orthovanadate, protease inhibitor cocktail (Roche), phosphatase inhibitor cocktail (Roche)) and then mixed with an equal part of 1% Triton X-100 (containing protease and phosphatase inhibitors). This mixture was incubated at 4°C with rotation for 10 minutes followed by centrifugation at 50,000x g for 20 minutes at 4°C to yield PSD preparation. PSD pellets were frozen and subsequently processed for mass spectrometry (see below).

### Mass spectrometry-based proteomics

The PSD pellet was reconstituted in 100 µl of lysis buffer (4% SDS in 0.1M Tris-HCl pH 7.6) and sonicated using a tip sonicator (Branson, SFX 550). Protein concentration was measured using BCA protein assays (Thermo, 23227). Proteins were reduced using 100 mM dithiothreitol for 30 min followed by alkylation with 50 mM iodoacetamide for 30 min. Around 50 µg of proteins were transferred to a 10 kDa centrifugal filter (Millipore Sigma, Amicon Ultra-0.5), in which the protein sample was mixed with 8 M urea (in 0.1 M triethylammonium bicarbonate, TEAB). The proteins on the membrane filter were centrifuged at 4,000 x g for 10 min to remove SDS followed by buffer exchange with 100 mM TEAB. Lys-C (FUJIFILM Wako, 125-05061) was added to the filter at an enzyme to protein ratio of 1: 50 and digested at 37 °C for overnight. Peptides were analyzed using Q Exactive HF mass spectrometer (Thermo Scientific, San Jose, CA) coupled to Ultimate 3000 liquid chromatography system (Thermo Scientific, San Jose, CA). Around 1 µg of peptides of individual samples were loaded onto a trap column (2 cm × 100 μm, Acclaim PepMap100, Thermo Fisher Scientific) and separated using 200 cm long micropillar array column (PharmaFluidics). The following gradient was generated with solvent A (0.1 % formic acid in water) and solvent B (0.1 % formic acid in acetonitrile): 5-7 % sol B in 15 min, 7-27 % sol B in 235 min, 27-40 % sol B in 15 min, 40-80 % sol B in 20 min, maintaining at 80 % sol B for 15 min, and equilibrating with 5 % sol B for 20 min. The flow rate was set as 300 nL min^−1^. Mass spectrometry data were acquired in data-dependent acquisition mode. Full MS scans were acquired for the mass range of 350-1800 m/z at the resolution of 120,000. Twenty most abundant ions were isolated under 1.6 *m/z* isolation width and fragmented with normalized collision energy (NCE) of 27. The MS/MS scans were acquired at a resolution of 30,000 with a fixed first m/z of 110 m/z. Maximum ion injection time was 50 ms and 54 ms for full MS and MS/MS scan, respectively. The raw mass spectrometry data were searched using Andromeda in MaxQuant software suite (version 2.0.1.0) against a UniProt mouse protein database (17,015 entries) along with common contaminants. Peptides with a maximum of two missed cleavages were considered. Acetylation of protein N-terminus, oxidation of methionine and deamidation of asparagine and glutamine were set as variable modifications and carbamidomethylation of cysteine was used as a fixed modification. False discovery rate was applied as 1% for both peptide and protein level. Quantification of SILAC pairs was performed with standard settings in MaxQuant.

The mass spectrometry proteomics data have been deposited to the ProteomeXchange Consortium via the PRIDE(Perez-Riverol et al., 2022) partner repository with the dataset identifier PXD065808.

### Mass spectrometry data analysis

The matrix of relative intensities of identified peptides from MaxQuant was processed using Python (SciPy v1.13.1, NumPy v1.26.4). For each peptide, the relative intensities corresponding to light (Lys-^12^C_6_, L) and heavy (Lys-^13^C_6_, H) lysine isotopes were extracted, and the relative isotope abundance (RIA) was calculated for each individual protein using the formula:

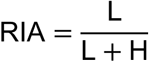

RIA values were determined for each protein across four experimental timepoints (0 days, 1 month, 3 months, and 6 months), with biological replicates consisting of 4 males and 3 females at day 0 (n = 7) and 3 males and 3 females at each subsequent timepoint (n = 6). For each protein, the RIA at day 0, denoted as 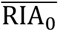 was calculated from these initial measurements, and protein turnover rate (t_1/2_) were subsequently estimated by fitting the RIA values for each protein to a single exponential decay model. This model employed the pre-calculated 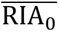 as a fixed initial abundance instead of 100%, with the best-fit protein turnover rate determined via non-linear least squares regression (implemented using SciPy’s “curve_fit” function), according to the following equation:

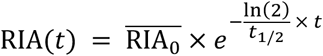

Following model fitting, proteins and their corresponding estimated half-lives were selected for downstream analysis only if they satisfied a set of stringent data and fit quality criteria: a coefficient of determination between fitted curve and the observed data of at least 0.8 (R² ≥ 0.8), average day0 RIA greather than 80% (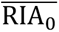 ≥ 0.8), no more than three missing values across all timepoints, and at least one valid RIA measurement at each timepoint.

Further analyses were conducted to compare protein half-lives both between sexes and between behaviorally manipulated experimental groups. For the sex-specific comparison, the primary dataset described in previous step was stratified into male and female cohorts. These cohorts were then analyzed separately, applying the identical analytical model, fitting strategies, and inclusion criteria used for the primary analysis. Samples for these sex-specific analyses were collected at 0 weeks (n = 4 for males, n = 3 for females), and at 1 month, 3 months, and 6 months (n = 3 for both males and females at each of these subsequent timepoints).

For comparisons involving behavioral manipulation, fear-conditioned (FC) male mice were compared to control male mice. Samples for this analysis were collected at 0 weeks (n = 4 for both FC and control groups), and at 1 week, 1 month, and 3 months (n = 3 for both groups at these subsequent timepoints). Crucially, for all these group-specific comparisons, protein half-lives were estimated employing the identical analytical formula (the single exponential decay model described previously), non-linear least squares regression fitting strategy, and all associated data and fit quality inclusion criteria as detailed for the primary analysis.

For select validation experiments, a modified double exponential decay function incorporating heavy amino acid recycling constants was applied to refine lifetime estimations (Fornasiero et al., 2018). This model accounts for two distinct recycled pools and was adapted from published formulations describing amino acid turnover in vivo:

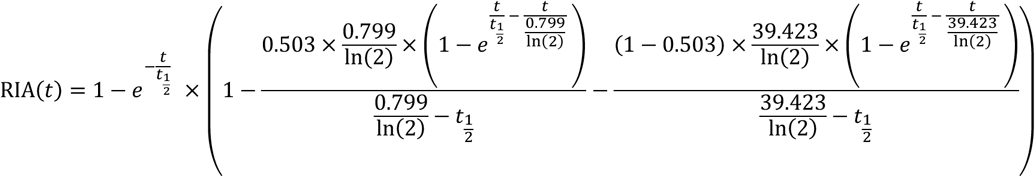

### Pathway analysis

All proteins passing filtering criteria in the combined dataset (n = 1526) were annotated via automated API calls to the Universal Protein Resource (UniProt Consortium) database hosted by EMBL-EBI (Nightingale et al., 2017). For each protein, associated Ensembl gene identifiers, sequence data, functional domains, and Gene Ontology (GO) annotations were retrieved. The Uniprot and Ensembl identifiers were used to cross-reference additional datasets, including the SFARI Gene database for autism spectrum disorder (ASD) risk genes (Abrahams et al., 2013). Human ASD risk genes were retrieved, and their associated risk scores were matched with our half-life dataset. Genes were considered ASD risk genes if they had an ASD risk score of 1-3 or were classified as syndromic genes.

To assess functional enrichment and protein-protein interactions, STRING network analysis was performed with custom background genes of all the proteins identified in mass spectrometry (Szklarczyk et al., 2023). The top 50 most stable proteins were subjected to unsupervised clustering using k-means (k = 6) to identify cluster labels, and significantly enriched non-redundant gene ontology terms are extracted. STRING analyses were also carried out on proteins exhibiting sex-specific differences in stability (defined by female-to-male half-life ratios exceeding ±2 standard deviations from the overall mean) and proteins with tuned stability after aversive experience (defined by cFC/control half-life ratios exceeding ±2 standard deviations from the mean), each clustered using k-means (k = 8). All STRING analyses significance of interaction networks was determined based on default STRING enrichment thresholds.

## Acknowledgements

NIH Grant support: R01MH112152 to R.L.H.; R00MH124920 to A.M.B.; U01CA271410 and P30CA15083 to A.P. We are grateful to the Huganir, Pandey and Bygrave lab members for their support and critical feedback on the project, particularly Ashley Irving, Sarah Rodriguez, and Richard Johnson for their technical assistance.

## Author contributions

S.H., D.M, A.P., A.M.B and R.L.H. designed research; S.H., D.M., and A.M.B. performed research; S.H., S.Z., D.M. and A.M.B. analyzed data; and A.M.B, S.Z., D.M., S.H., and R.L.H. wrote the paper.

## Supplementary figures

**Supplementary Figure 1.**
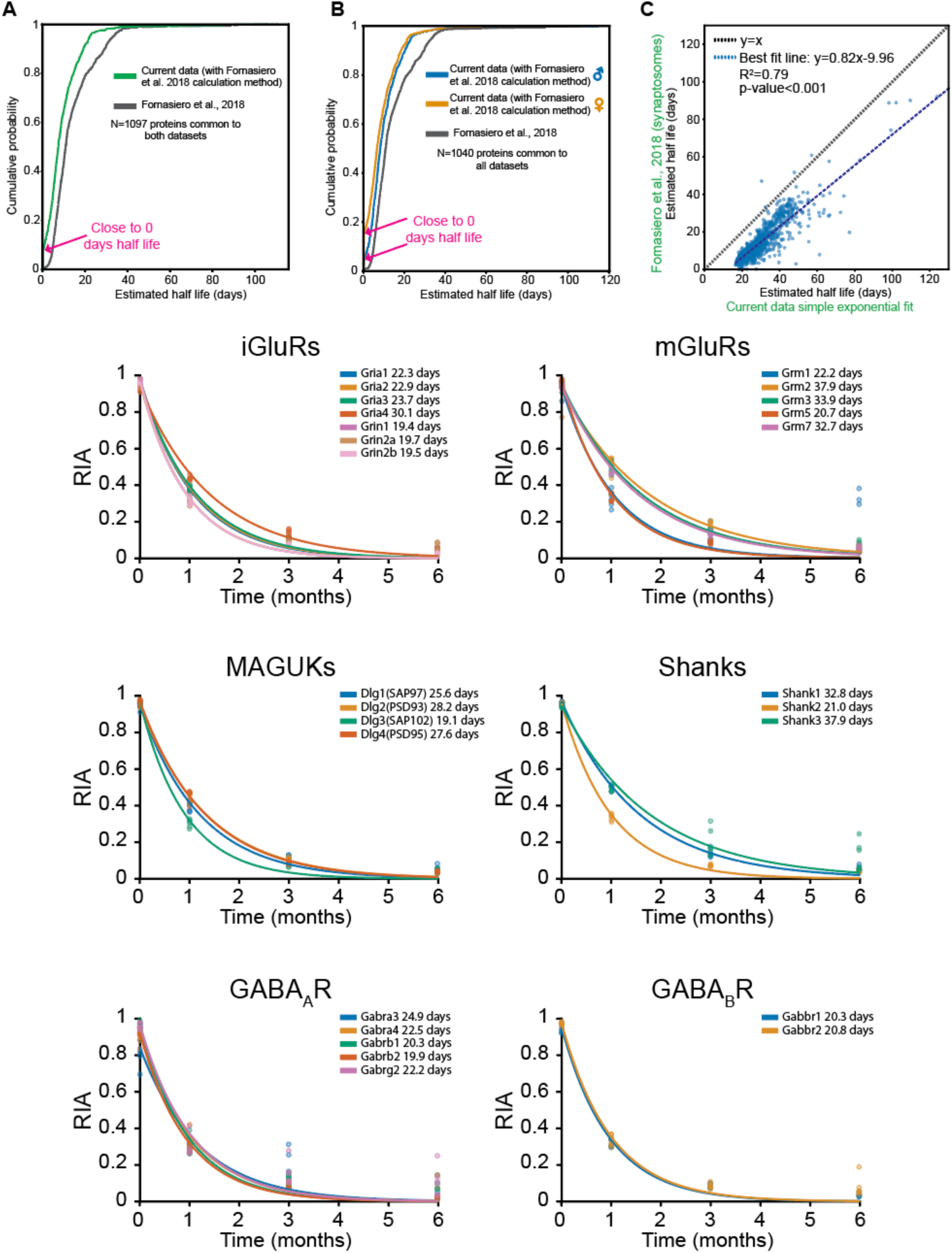
Calculation of relative protein turnover rates. (**A**) Cumulative probability plot showing our data (green) compared to the data from Fornasiero *et al*. (gray) for the 1097 common protein using the constants from Fornasiero *et al.,* 2018. Note the cluster of protein with estimates half-lives close to 0 days. (**B**) Same as (A) except split by sex with data from male animals (blue) and female animals (orange) plotted separately for the 1040 common proteins between the datasets. Note that the female animals in our dataset had a more substantial proportion of proteins with half-lives close to zero. (**C**) Scatter plot of common protein half-lives (N=1117) our simple exponential method plotted on the x-axis and half-lives reported by Fornasiero *et al*. on the y-axis. Reassuringly, there was a strong and significant positive correlation observed (Spearman’s Rank Correlation p-value< 0.001). Note that there was an overall trend to our method overestimating the estimated protein half-lives. (**D-I**) Example data showing the RIA decay for different protein family members. Each dot represents data from a single animal and the line shows the exponential fit that was used to estimate the protein half-life.

**Supplementary Figure 2.**
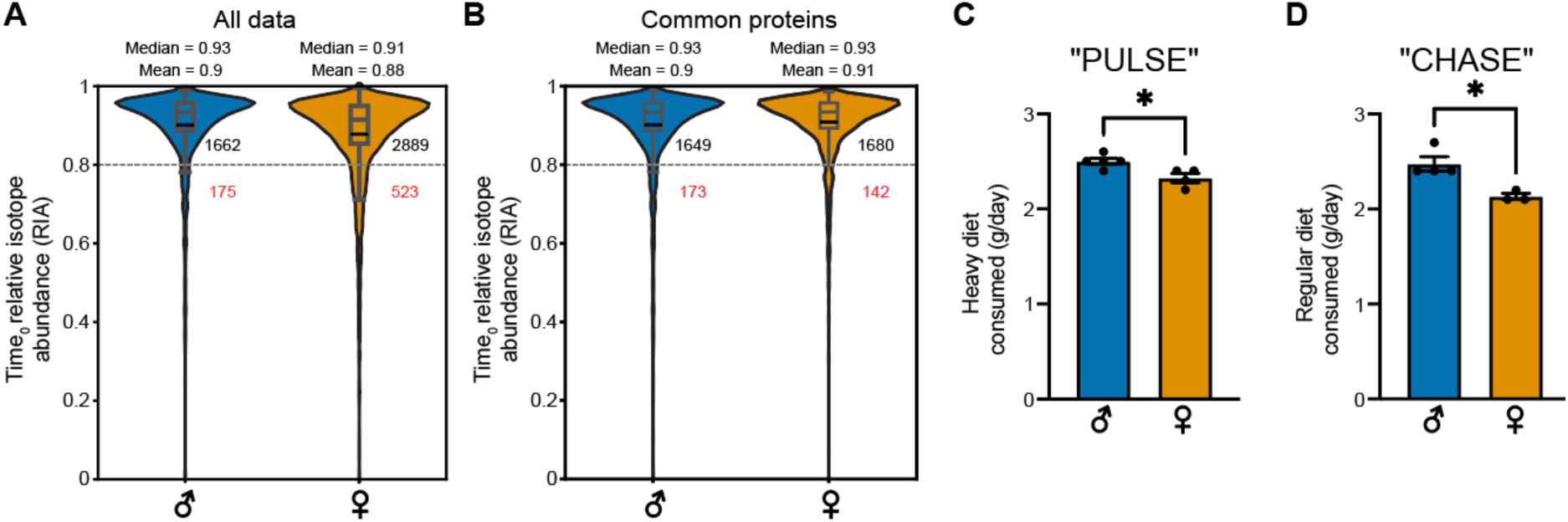
Sex differences in food intake. (**A**) Violin plot showing the starting (time 0) RIA values for all proteins split by sex. The median and mean values are indicated, along with the number of proteins below the 0.8 threshold (which were not included for subsequent analyses). (**B**) Same as in (A) except filtered to only display common proteins found in both male and female animals. (**C, D**) Average diet consumed per mouse per day during the labeling (C) and chase (D) phases. Note, as animals were group housed each data point represents data from one cage. Bars display mean and error bards display standard error of the mean. In general, female animals consumed less food than males.

**Supplementary Figure 3.**
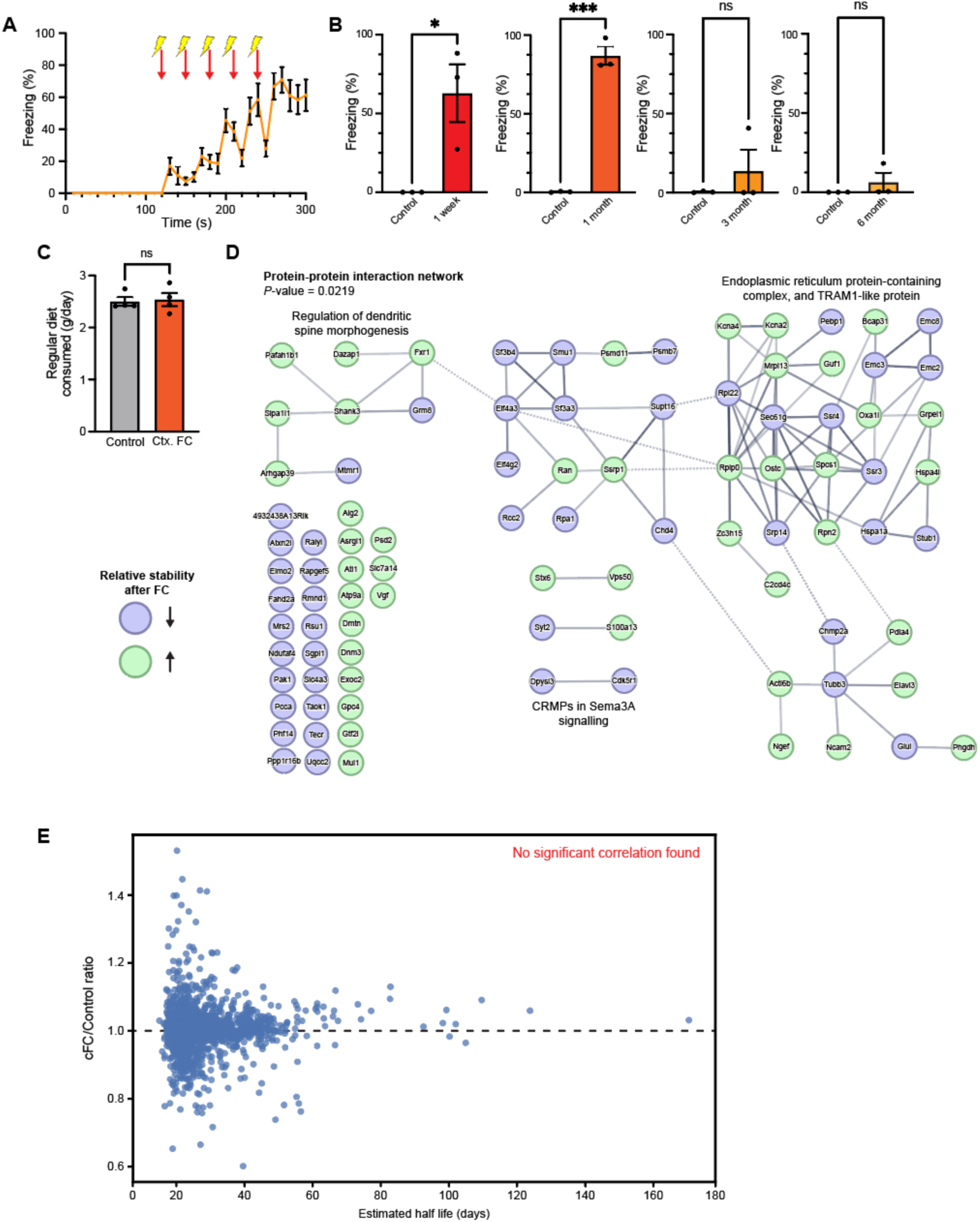
Experience dependent changes in protein stability. (**A**) Freezing levels during the conditioning session of labelled mice. The freezing levels robustly increased after delivery of the electric foot shocks. (**B**) Freezing levels of the animals in the 1 week, 1 month, and 3-month groups immediately before tissue was collected. (**C**) There was no difference in the amount of food consumed in the control and cFC groups. (**D**) STRING analysis of the proteins with +/− 2 SD differences in turnover rates between the control and cFC groups. Green shading indicates greater stability following cFC and blue shading indicates decreased stability following cFC. (**E**) Scatter plot showing no correlation between the overall protein stability (x-axis) and the ratio of cFC/control turnover rates. Throughout, bars or lines displays the mean, and the error bars the standard error of the mean.

**Supplementary Table 1.**
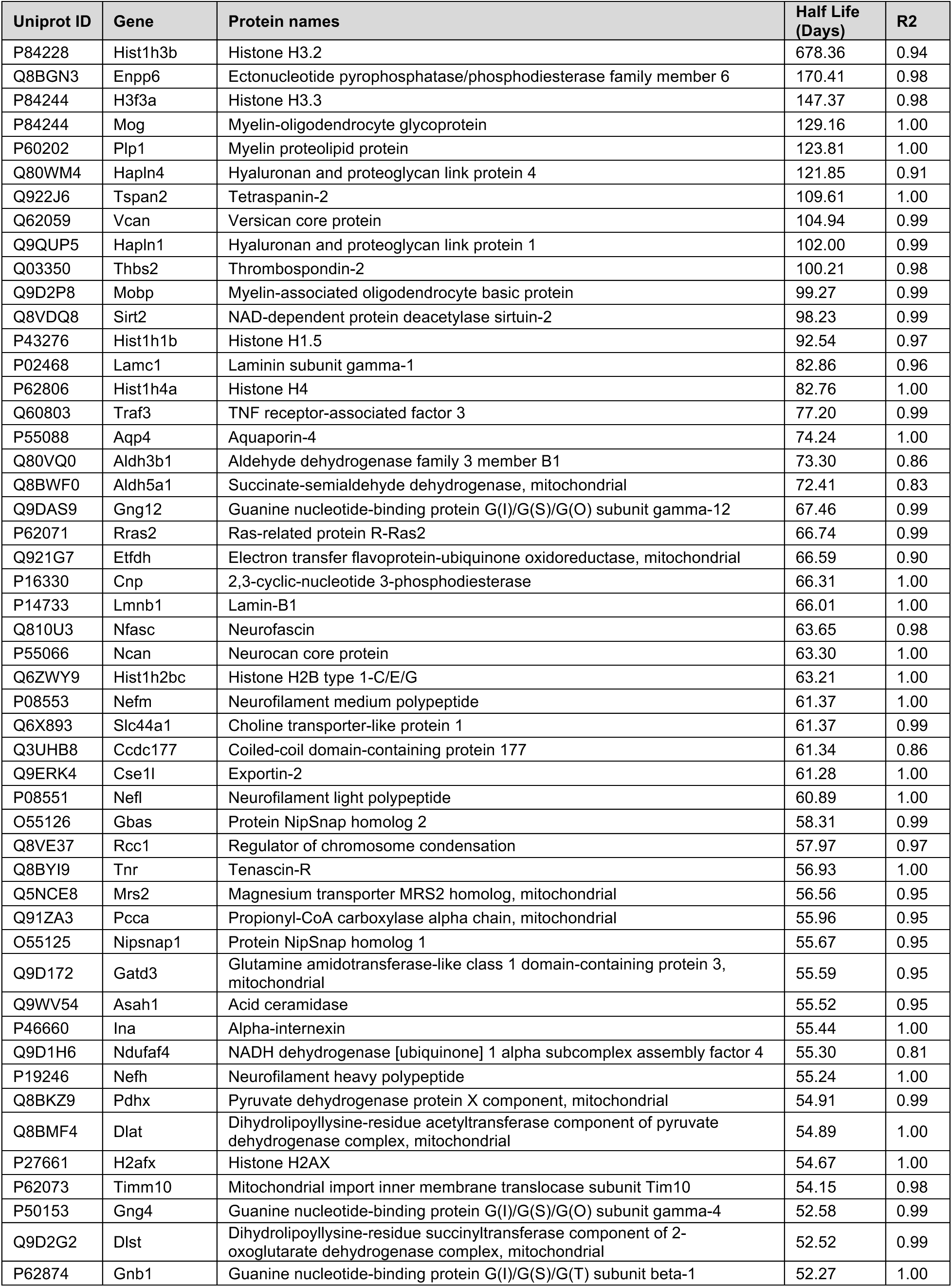
Top 50 most stable proteins combining male and female.

**Supplementary Table 2.**
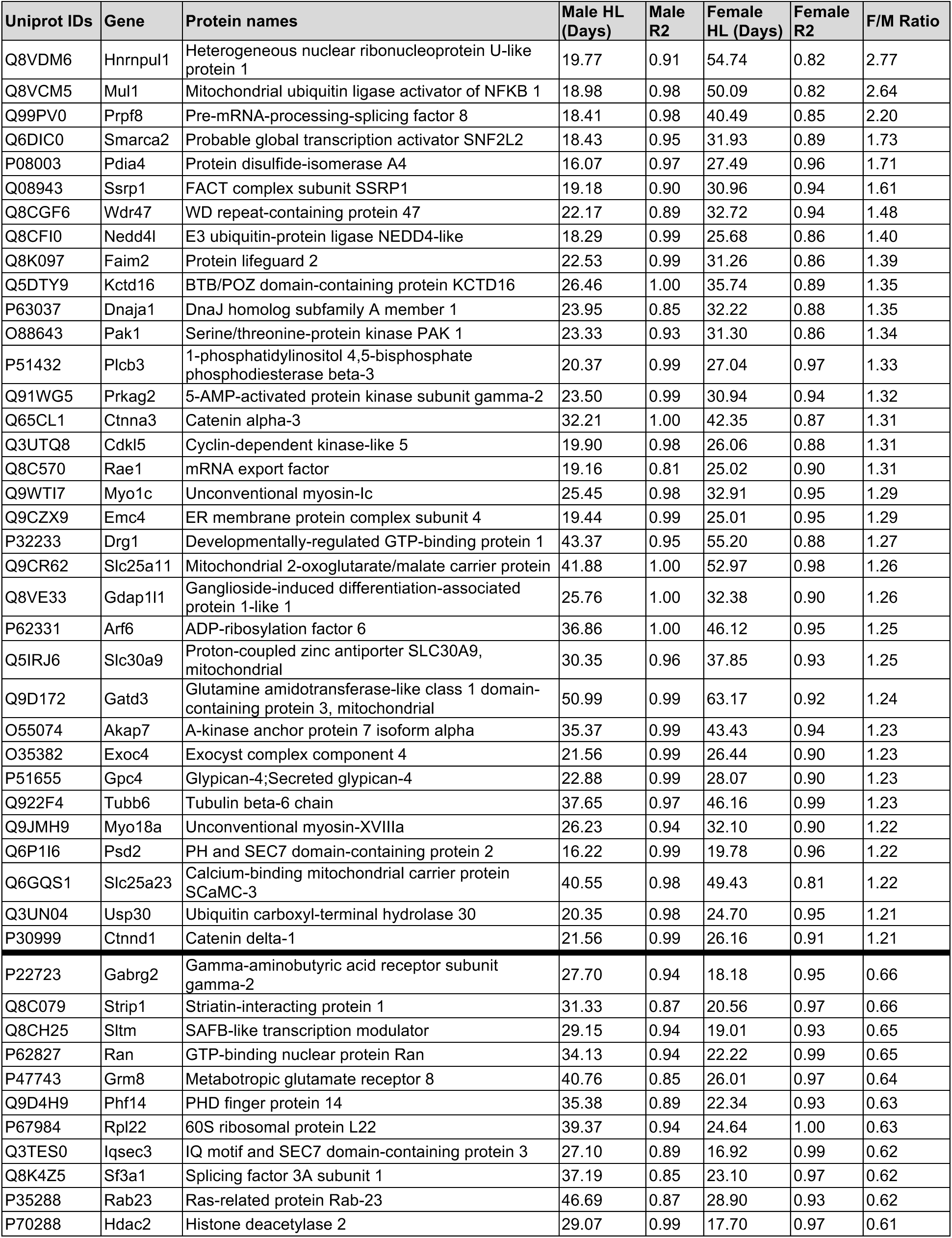
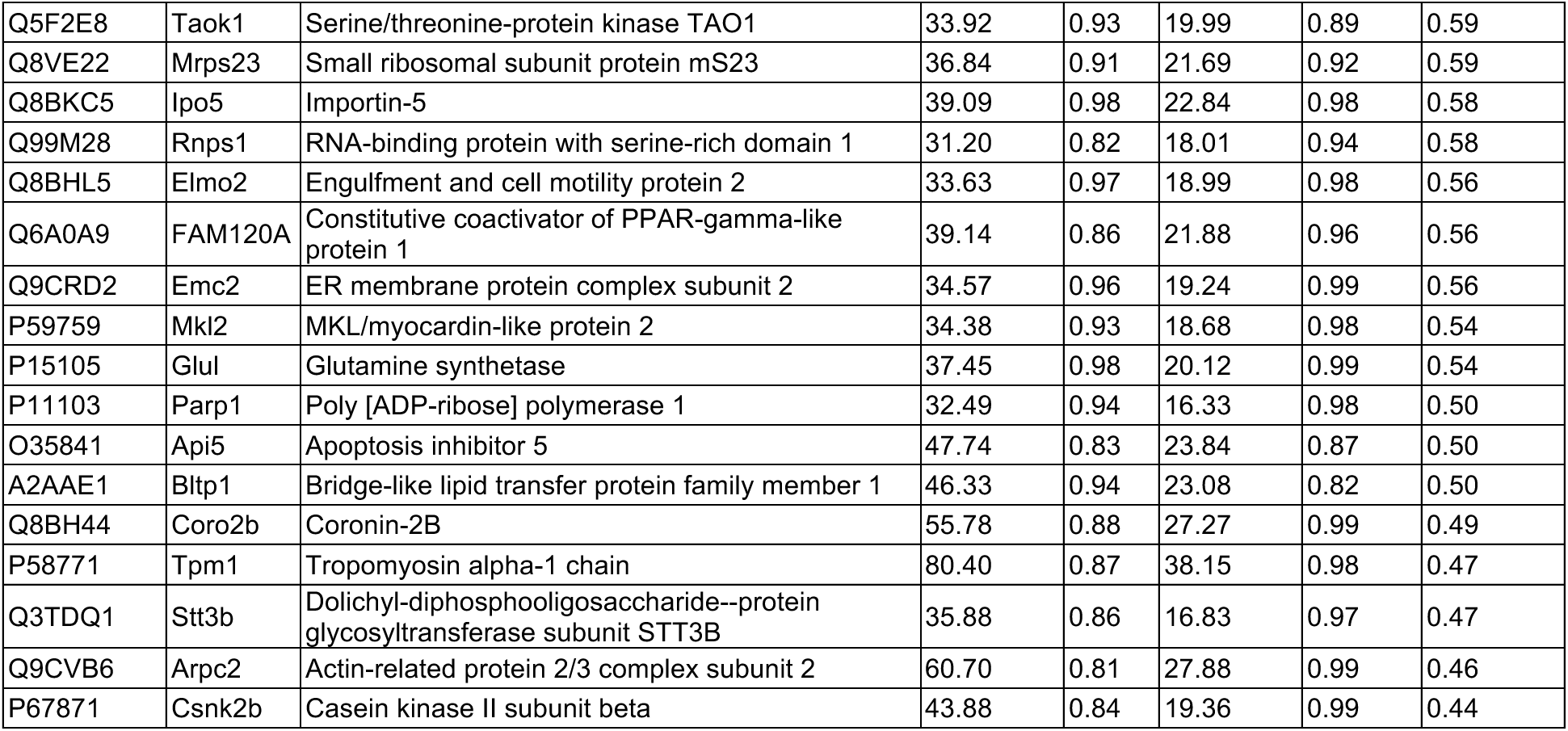
Proteins with sex stability difference +/− 2 standard deviations (SD) from the mean female to male ratio.

**Supplementary Table 3.**
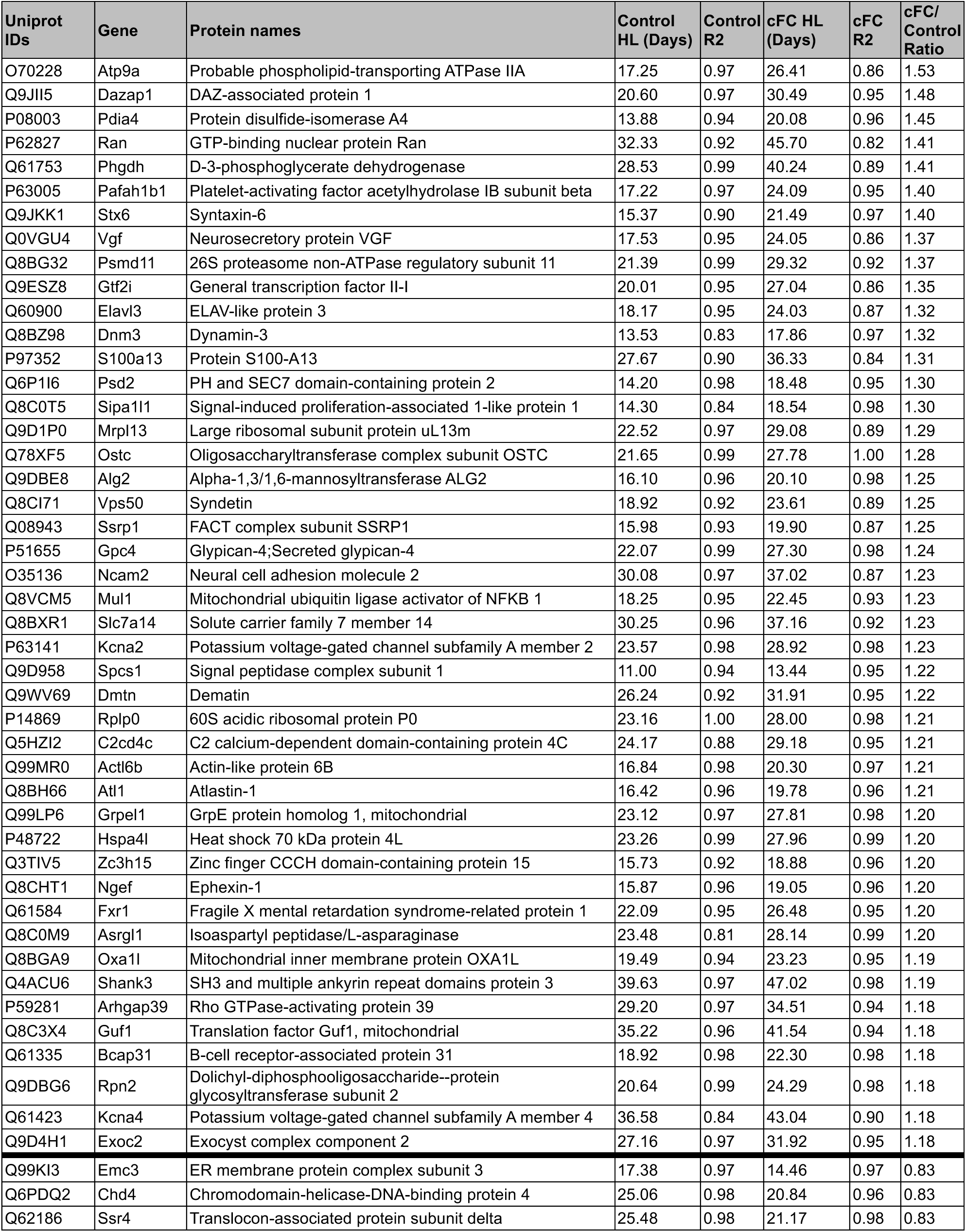
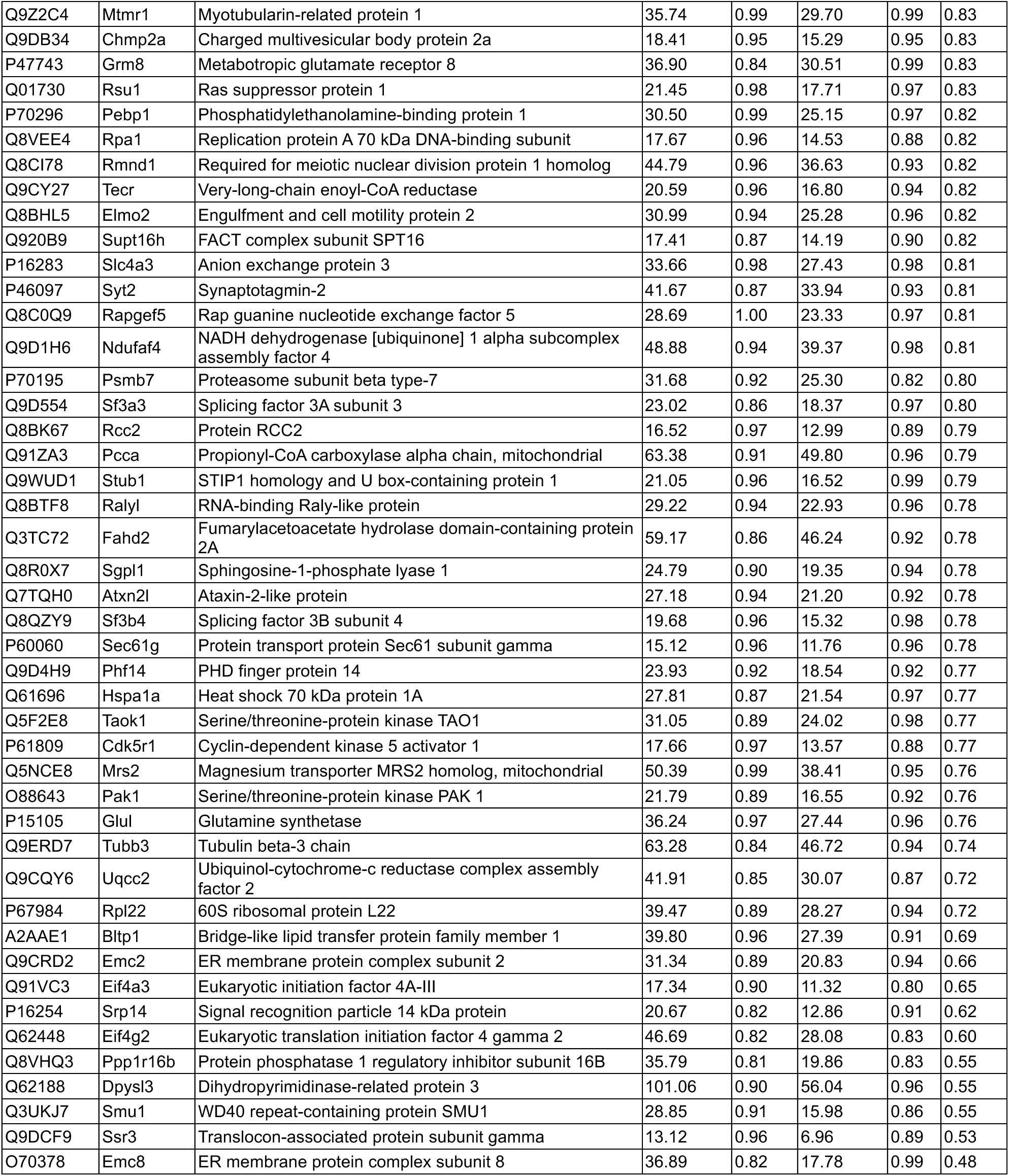
Proteins with aversive experience stability difference +/− 2 standard deviations from the fear condition to control ratio in male mice.

**Supplementary Table 4:**
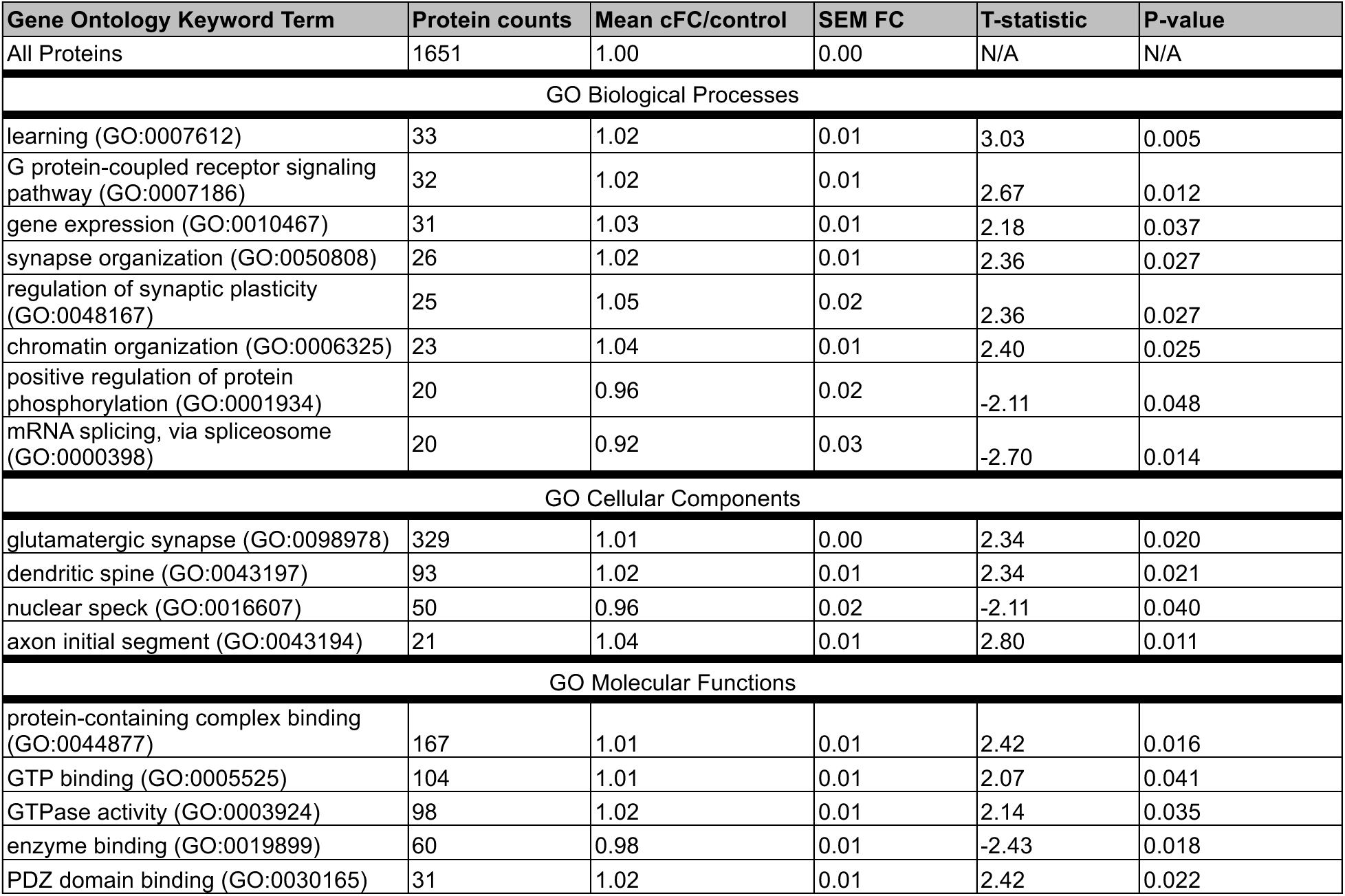

## References

Abrahams BS, Arking DE, Campbell DB, Mefford HC, Morrow EM, Weiss LA, Menashe I, Wadkins T, Banerjee-Basu S, Packer A (2013) SFARI Gene 2.0: A community-driven knowledgebase for the autism spectrum disorders (ASDs). Mol Autism 4.

Bonkhoff AK, Coughlan G, Perosa V, Alhadid K, Schirmer MD, Regenhardt RW, Van Veluw S, Buckley R, Fox MD, Rost NS (2025) Sex differences in age-associated neurological diseases-A roadmap for reliable and high-yield research. Available at: https://www.science.org.

Bulovaite E, Qiu Z, Kratschke M, Zgraj A, Fricker DG, Tuck EJ, Gokhale R, Koniaris B, Jami SA, Merino-Serrais P, Husi E, Mendive-Tapia L, Vendrell M, O’Dell TJ, DeFelipe J, Komiyama NH, Holtmaat A, Fransén E, Grant SGN (2022) A brain atlas of synapse protein lifetime across the mouse lifespan. Neuron 110:4057–4073.e8.

Bygrave AM, Sengupta A, Jackert EP, Ahmed M, Adenuga B, Nelson E, Goldschmidt HL, Johnson RC, Zhong H, Yeh FL, Sheng M, Huganir RL (2023) Btbd11 supports cell-type-specific synaptic function. Cell Rep 42.

Cyranowski JM, Frank E, Young E, Shear; M Katherine (2000) Adolescent Onset of the Gender Difference in Lifetime Rates of Major Depression A Theoretical Model.

Dankovich TM, Rizzoli SO (2022) The Synaptic Extracellular Matrix: Long-Lived, Stable, and Still Remarkably Dynamic. Front Synaptic Neurosci 14.

Dörrbaum AR, Kochen L, Langer JD, Schuman EM (2018) Local and global influences on protein turnover in neurons and glia. Elife 7 Available at: http://www.ncbi.nlm.nih.gov/pubmed/29914620.

Fernández E, Collins MO, Uren RT, Kopanitsa M V., Komiyama NH, Croning MDR, Zografos L, Armstrong JD, Choudhary JS, Grant SGN (2009) Targeted tandem affinity purification of PSD-95 recovers core postsynaptic complexes and schizophrenia susceptibility proteins. Mol Syst Biol 5.

Fornasiero EF et al. (2018) Precisely measured protein lifetimes in the mouse brain reveal differences across tissues and subcellular fractions. Nat Commun 9.

Frey U, Morris RG (1998) Synaptic tagging: implications for late maintenance of hippocampal long-term potentiation. Trends Neurosci 21:181–188 Available at: http://www.ncbi.nlm.nih.gov/pubmed/9610879.

Geiger JR, Lübke J, Roth A, Frotscher M, Jonas P (1997) Submillisecond AMPA receptor-mediated signaling at a principal neuron-interneuron synapse. Neuron 18:1009–1023 Available at: http://www.ncbi.nlm.nih.gov/pubmed/9208867.

Geiger JR, Melcher T, Koh DS, Sakmann B, Seeburg PH, Jonas P, Monyer H (1995) Relative abundance of subunit mRNAs determines gating and Ca2+ permeability of AMPA receptors in principal neurons and interneurons in rat CNS. Neuron 15:193–204 Available at: http://www.ncbi.nlm.nih.gov/pubmed/7619522.

Heo S, Diering GH, Na CH, Nirujogi RS, Bachman JL, Pandey A, Huganir RL (2018) Identification of long-lived synaptic proteins by proteomic analysis of synaptosome protein turnover. Proc Natl Acad Sci U S A 115:E3827–E3836 Available at: http://www.ncbi.nlm.nih.gov/pubmed/29610302.

Kimbrel NA et al. (2023) Identification of Novel, Replicable Genetic Risk Loci for Suicidal Thoughts and Behaviors Among US Military Veterans. JAMA Psychiatry 80:135–145.

Levey DF, Stein MB, Wendt FR, Pathak GA, Zhou H, Aslan M, Quaden R, Harrington KM, Nuñez YZ, Overstreet C, Radhakrishnan K, Sanacora G, McIntosh AM, Shi J, Shringarpure SS, Concato J, Polimanti R, Gelernter J (2021) Bi-ancestral depression GWAS in the Million Veteran Program and meta-analysis in >1.2 million individuals highlight new therapeutic directions. Nat Neurosci.

Li W, Dasgupta A, Yang K, Wang S, Hemandhar-Kumar N, Chepyala SR, Yarbro JM, Hu Z, Salovska B, Fornasiero EF, Peng J, Liu Y (2025) Turnover atlas of proteome and phosphoproteome across mouse tissues and brain regions. Cell Available at: http://www.ncbi.nlm.nih.gov/pubmed/40118046.

Loomes R, Hull L, Mandy WPL (2017) What Is the Male-to-Female Ratio in Autism Spectrum Disorder? A Systematic Review and Meta-Analysis. J Am Acad Child Adolesc Psychiatry 56:466–474.

Melander JB, Nayebi A, Jongbloets BC, Fortin DA, Qin M, Ganguli S, Mao T, Zhong H (2021) Distinct in vivo dynamics of excitatory synapses onto cortical pyramidal neurons and parvalbumin-positive interneurons. Cell Rep 37.

Mohar B et al. (2025) DELTA: a method for brain-wide measurement of synaptic protein turnover reveals localized plasticity during learning. Nat Neurosci Available at: http://www.ncbi.nlm.nih.gov/pubmed/40164741.

Nievergelt CM et al. (2024) Genome-wide association analyses identify 95 risk loci and provide insights into the neurobiology of post-traumatic stress disorder. Nat Genet 56:792–808.

Nightingale A, Antunes R, Alpi E, Bursteinas B, Gonzales L, Liu W, Luo J, Qi G, Turner E, Martin M (2017) The Proteins API: accessing key integrated protein and genome information. Nucleic Acids Res 45:W539–W544 Available at: http://www.ncbi.nlm.nih.gov/pubmed/28383659.

Pelkey KA, Barksdale E, Craig MT, Yuan X, Sukumaran M, Vargish GA, Mitchell RM, Wyeth MS, Petralia RS, Chittajallu R, Karlsson RM, Cameron HA, Murata Y, Colonnese MT, Worley PF, McBain CJ (2015) Pentraxins coordinate excitatory synapse maturation and circuit integration of parvalbumin interneurons. Neuron 85:1257–1272.

Perez-Riverol Y, Bai J, Bandla C, García-Seisdedos D, Hewapathirana S, Kamatchinathan S, Kundu DJ, Prakash A, Frericks-Zipper A, Eisenacher M, Walzer M, Wang S, Brazma A, Vizcaíno JA (2022) The PRIDE database resources in 2022: A hub for mass spectrometry-based proteomics evidences. Nucleic Acids Res 50:D543–D552.

Savas JN, Toyama BH, Xu T, Yates JR, Hetzer MW (2012) Extremely long-lived nuclear pore proteins in the rat brain. Science (1979) 335:942.

Szklarczyk D, Kirsch R, Koutrouli M, Nastou K, Mehryary F, Hachilif R, Gable AL, Fang T, Doncheva NT, Pyysalo S, Bork P, Jensen LJ, Von Mering C (2023) The STRING database in 2023: protein-protein association networks and functional enrichment analyses for any sequenced genome of interest. Nucleic Acids Res 51:D638–D646.

Trachtenberg JT, Chen BE, Knott GW, Feng G, Sanes JR, Welker E, Svoboda K (2002) Long-term in vivo imaging of experience-dependent synaptic plasticity in adult cortex. Available at: www.nature.com/nature.

Tsien RY (2013) Very long-term memories may be stored in the pattern of holes in the perineuronal net. Proc Natl Acad Sci U S A 110:12456–12461.

Wang X, McCoy PA, Rodriguiz RM, Pan Y, Je HS, Roberts AC, Kim CJ, Berrios J, Colvin JS, Bousquet-Moore D, Lorenzo I, Wu G, Weinberg RJ, Ehlers MD, Philpot BD, Beaudet AL, Wetsel WC, Jiang YH (2011) Synaptic dysfunction and abnormal behaviors in mice lacking major isoforms of Shank3. Hum Mol Genet 20:3093–3108.

Yang Y, Fang F, Arnberg FK, Kuja-Halkola R, D’Onofrio BM, Larsson H, Brikell I, Chang Z, Andreassen OA, Lichtenstein P, Valdimarsdóttir UA, Lu D (2024) Sex differences in clinically diagnosed psychiatric disorders over the lifespan: a nationwide register-based study in Sweden. The Lancet Regional Health - Europe.

Zheng CY, Petralia RS, Wang YX, Kachar B, Wenthold RJ (2010) Sap102 is a highly mobile maguk in spines. Journal of Neuroscience 30:4757–4766.

